# *AGL16* regulates genome-wide gene expression and flowering time with partial dependency on *SOC1* in *Arabidopsis*

**DOI:** 10.1101/2021.05.10.443448

**Authors:** Xue Dong, Li-Ping Zhang, Dong-Mei Yu, Fang Cheng, Yin-Xin Dong, Xiao-Dong Jiang, Fu-Ming Qian, Franziska Turck, Jin-Yong Hu

## Abstract

Flowering transition is pivotal and tightly regulated by complex gene-regulatory-networks, in which AGL16 plays important roles. But the molecular function and binding property of AGL16 is not fully explored *in vivo*. With ChIP-seq and comparative transcriptomics approaches, we characterized the AGL16 targets spectrum and tested its close molecular and genetic interactions with *SOC1*, the key flowering integrator. AGL16 bound to promoters of more than 2000 genes via CArG-box motifs that were highly similar to that of SOC1. Being consistent with this, AGL16 formed protein complex and shared a common set of targets with SOC1. However, only very few genes showed differential expression in the *agl16-1* loss-of-function mutant, whereas in the *soc1-2* knockout background, AGL16 repressed and activated the expression of 375 and 182 genes, respectively, with more than a quarter of the DEGs were also bound by AGL16. AGL16 targeted potentially to about seventy flowering time genes involved in multiple pathways. Corroborating with these, *AGL16* repressed the flowering time stronger in *soc1-2* than in Col-0 background. These data reveals that AGL16 regulates gene expression and flowering time with a partial dependency on *SOC1* activity. Moreover, AGL16 participated in the regulation of water loss and seed dormancy. Our study thus defines the AGL16 molecular spectrum and provides insights underlining the molecular coordination of flowering and environmental adaptation.

## Introduction

Timely transitions from vegetative to reproductive growth (floral transition) and from dormant to germinating seeds (dormancy release) determine the capacity of plants to adapt to changing environments, thus these processes are under tight control by complex interactions between endogenous signals and exogenous environmental factors (Andres and Coupland 2012; Michaels 2009; Nee, Xiang, and Soppe 2017). The gene-regulatory-network (GRN) controlling floral transition converges at several floral integrator genes like *SUPPRESSOR OF CONSTANS 1* (*SOC1*) and *FLOWERING LOCUS T* (*FT*). These genes often encode transcription regulators controlling the transcription of their downstream targets by binding to specific *cis*-motifs, for example CArG-boxes (Andres and Coupland 2012; Michaels 2009; Fornara, de Montaigu, and Coupland 2010). CArG-box motifs are binding sites specific for MADS-box transcription factors (TFs) like SOC1, FLOWEING LOCUS C (FLC), SHORT VEGETATIVE PHASE (SVP) and SEPALLATA 3 (SEP3) (Gregis et al. 2013; Mateos et al. 2015; Mateos et al. 2017; Kaufmann et al. 2009; Deng et al. 2011; Immink et al. 2009; Immink et al. 2012; Kaufmann et al. 2010; Tao et al. 2012; Aerts et al. 2018). These MADS-box TFs often form homo- and/or hetero-protein complexes that act in concert and bind to the CArG-box motifs in promoters of more than hundreds of downstream genes to regulate flowering time and other developmental processes of *Arabidopsis thaliana*.

SOC1 is one key flowering promoter integrating signals from photoperiod, temperature, hormones and age-related pathways (Lee and Lee 2010). SOC1 forms protein complex with AGL24 to activate *LFY* and *AP1* to initiate and maintain flower meristem identity but represses *SEP3* to prevent premature differentiation of floral meristem (Lee et al. 2008). SOC1 can promote the expression of *TARGET OF FLC AND SVP1* (*TFS1*) via recruiting histone demethylase RELATED TO EARLY FLOWERING 6 (REF6) and chromatin remodeler BRAHMA (BRM), and cooperates with SQUAMOSAL PROMOTER BINDING PROTEIN-LIKE 15 (SPL15) to modulate their targets expression thereby regulating flowering time (Richter et al. 2019; Hyun et al. 2016). SOC1 forms a set of heterologous complexes with other MADS-box transcription factors, for example AGL16 (de Folter et al. 2005; Immink et al. 2009). Furthermore, *SOC1* times flowering downstream of several hormone signaling pathways including GA, ABA and BRs (Hwang et al. 2019; Jung et al. 2012; Li et al. 2017) and of nutrient status (Yan et al. 2021; Olas et al. 2019; Liu et al. 2013). Interestingly, profiling of SOC1 targets also identifies genes involved in the signaling processes of these hormones and nutrients (Immink et al. 2012; Tao et al. 2012). However, the biological significance of these molecular interactions remains to be explored further.

AGL16 represses flowering with dependency on the genetic background, the photoperiod of growth conditions, and gene dosages in *A. thaliana* (Hu et al. 2014). Only under the inductive long-day conditions loss-of-function mutants for *AGL16* show early flowering especially in the functional *FRI-FLC* background (Johanson et al. 2000; Michaels and Amasino 2001; Hu et al. 2014). *AGL16* expression can be modulated by the level of the Brassicaceae-specific *miR824*, for which natural variation has been reported (Hu et al. 2014; Kutter et al. 2007; de Meaux et al. 2008; Fahlgren et al. 2007; Rajagopalan et al. 2006). Interestingly, changes in *miR824* expression result in a significant modification of the plant flowering (Hu et al. 2014). *AGL16* acts in flowering time regulation via transcriptional regulation of *FT*, whose expression is also regulated by other MADS-box repressors such as SVP and FLC and other TFs (Aukerman and Sakai 2003; Searle et al. 2006; Jung et al. 2007; Castillejo and Pelaz 2008; Li et al. 2008; Mathieu et al. 2009). AGL16 forms complexes with SVP and FLC, and mildly represses their expression (Hu et al. 2014). *AGL16* is a direct downstream target of both FLC and SVP, but the expression of *AGL16* changes only weakly in loss-of-function mutants of both genes (Deng et al. 2011; Gregis et al. 2013; Mateos et al. 2015). Yeast-two-hybrids assays suggest that AGL16 interacts with SOC1 and other MADS-box TFs and it has been hypothesized that AGL16 could modulate the *SOC1* expression (de Folter et al. 2005; Immink et al. 2012; Immink et al. 2009). However, the exact AGL16 target spectrum and the impact of interactions between AGL16 and its partners remain under-explored.

In this study, we examined the molecular profiles that AGL16 bound and tested the molecular and genetic interactions between *AGL16* and *SOC1*. We found that, in contrast to its mild effects in flowering time regulation in Col-0 background, AGL16 could in fact bind to more than 2000 target genes that were involved in regulation of flowering time and other biological processes. We confirmed the molecular and genetic interactions of AGL16 with SOC1 and found that they shared many common targets. We demonstrated that the regulatory roles of AGL16 on genome-wide gene expression and flowering time depended partially on the SOC1 activity.

## Results

### AGL16 binds to a large set of genomic segments with CArG boxes

We profiled AGL16 binding sites by a ChIP-seq approach (chromatin immuno-precipitation followed by sequencing). We used a line expressing *AGL16* fused to a combined Yellow Fluorescent Protein (YFP)-HA epitope tag under the control of the Cauliflower Mosaic Virus 35S promoter (*AGL16OX*), which restores the early flowering of *agl16-1* to wild type Col-0 level (Fig. S1) (Hu et al. 2014). In two independent trials, we identified respectively 5463 and 3294 DNA segments statistically enriched for AGL16 binding, of which 3086 were shared (Table S2, S3). Most of the peaks were around 150-500 bp in both trials (Fig. S2). To test whether these segments were real binding sites for AGL16, we carried out ChIP-qPCR assays with two independent chromatin preparations for 20 peaks identified by ChIP-seq. These efforts confirmed 12 regions bound by AGL16-YFP-HA with a minimum two-fold enrichment in the *AGL16OX* line compared to *agl16-1* background (Fig. 1). Hence, a majority proportion of peaks detected via ChIP-seq method were reproducibly enriched.

**Fig 1.**
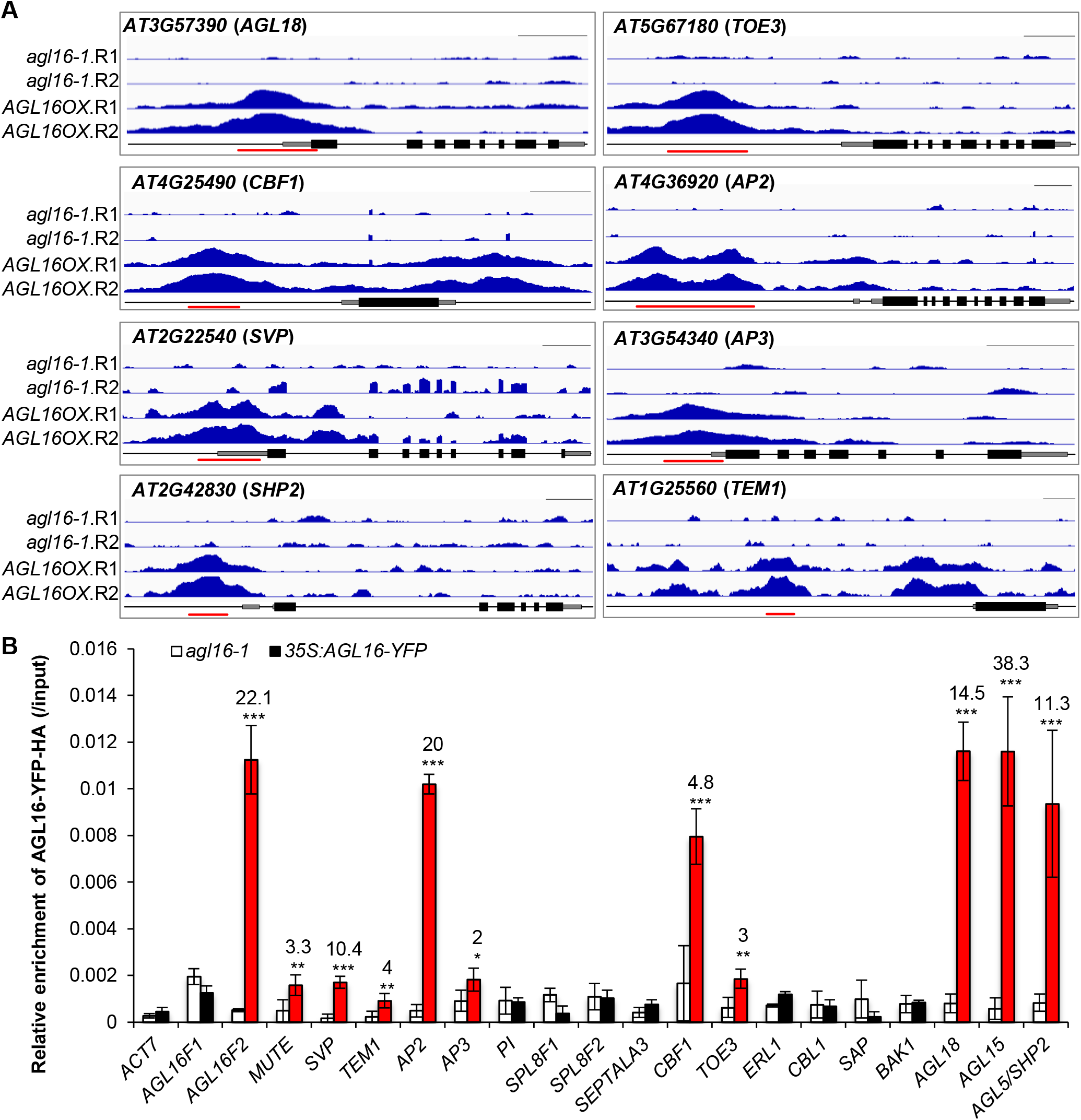
Validation of the AGL16 binding on target DNA fragments. **A**. Binding profiles for selected target genes. The TAIR10 annotation of the genomic locus was shown at the bottom of each box. For each panel, the profiles for two trials (R1 and R2) in *agl16-1* background line were shown in the upper panel, while the profiles for *agl16-1 35S:AGL16-YFP-HA* (*AGL16OX;* two trials) were shown in the middle panel of each box. All the genes were from 5’-end to 3’-end with scale bars indicating sequence lengths of 500 bp. Note that data range for each gene in *agl16-1* and *AGL16OX* was the same scale, but different genes could have different scale. Red lines marked the binding regions tested via ChIP-qPCR assays (**B**). **B**. ChIP-qPCR validation of AGL16 binding on 20 DNA segments. Significant enrichment (red bars) was defined with the following criteria: mean enrichment must be at least two-fold higher than negative control *ACT7*, the enrichment for *AGL16OX* (in *agl16-1* background) than *agl16-1* must be higher than two-fold change, and the amplification C_T_ number of IP samples must be at least 2 cycles less than no-antibody controls. This experiment was repeated with another independent trial, in which the relative enrichment of *AGL16F1* and *SAP* did not meet above criteria (see Figure S4). Statistics was carried out with *Student’s t-test* with Bonferroni correction. ***, P<0.001; **, p<0.01; *, p<0.05.

Peaks bound by AGL16 were annotated using Arabidopsis TAIR10 data to profile their distribution to genomic features (Fig. 2). The peaks from both trials were centered to the 3 Kb regions around transcriptional start sites (TSS; Fig. 2B). Around 60% of peaks located in the 1 Kb regions surrounding TSS (Fig. 2C; Table S3). About 10% of peaks were located in the 1-2 kb promoter regions upstream of TSS, while 10-12% of peaks were in exons/introns. Thus, AGL16 bound to DNA fragments close to TSS of a large set of genes. The 2339 genes with peaks mapped to gene body or up to 2 Kb upstream of their TSS were taken as AGL16 targets (Table S3).

**Fig 2.**
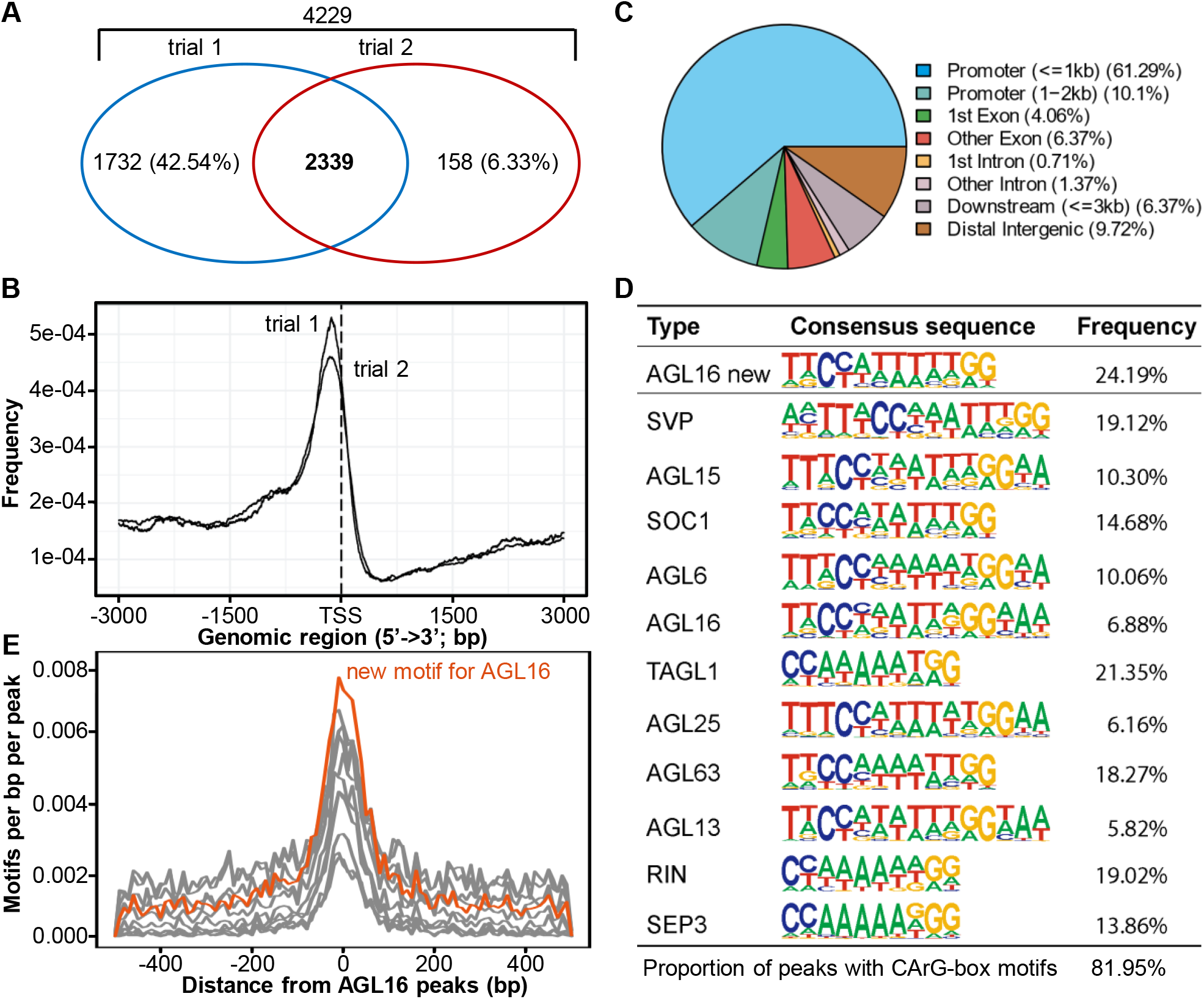
Genome-wide identification of AGL16 target genes via ChIP-seq. **A**. Venn diagram of AGL16 targets identified in two independent trials. **B**. Distribution of AGL16 binding sites for two trials surrounding the transcriptional starting site (TSS). **C**. Location distribution in relative to nearby genes for AGL16 binding sites of trial 1. Peaks within the 3 Kb promoter region were taken as AGL16 targets. **D**. CArG type of motifs over-represented in the AGL16 binding peaks. AGL16 new, which was highly similar to known SOC1 type, showed the *de novo* motif predicted for AGL16. Frequency gave the percentage for each motif presented in the binding peaks. **E**. Distribution of new (orange) and known (gray; shown in **D**) CArG type of motifs around AGL16 peaks center.

We next searched for potential cis-elements in the common peaks bound by AGL16 using HOMER, which could predict new motifs and identify known motifs (Heinz et al. 2010). This analysis reported a *de novo* CArG-box motif CCATTTTTGG for AGL16 in 707 peaks (24.2% of all common peaks; *Fisher P*=1e-340, in comparison to 3.8% at genome level; Fig. 2D, Table S3). Ten other CArG-box motifs were also significantly enriched, and matched to the known motifs of SVP, SOC1, SEP3, TAGL1, AGL63, and other MADS-box TFs, most of which could potentially interact with AGL16 (Fig. 2D; Fig. S3; Table S3). The *de novo* and the ten significantly enriched CArG-box motifs were all distributed around the peak center, indicating that AGL16 bound to its targets via the cluster of CArG-box motifs, just like SOC1 and other MADS-box proteins did (Tao et al. 2012; Immink et al. 2012; Deng et al. 2011). There were also other motifs significantly enriched in the AGL16 bound peaks, such as those bound by TCPs (321 peaks), bHLHs (1131), C2C2 DOFs (2524), WRKYs (1039). However, these motifs were not in the peaks center. Since AGL16 modulated significantly the flowering time in *Arabidopsis* (Hu et al. 2014), we next asked which flowering time genes could be targeted by AGL16.

### AGL16 targets flowering time genes in multiple pathways

The Arabidopsis genome contains ~400 flowering time genes, among which around 70 were targeted by AGL16 (Fig. 3; Table S3). This number was significantly larger than randomly expected (*Yates*’ *Chi-square test*, p<0.0001). Consistent with the described photoperiod dependency for *AGL16*-mediated flowering regulation (Hu et al. 2014), 37 genes (for example *AGAMOUS LIKE 15/16/18* (*AGL15/AGL16/AGL18*), *CONSTANS LIKE 1/3/4/5* (*COL1/3/4/5*), *TWIN SISTER OF FT* (*TSF*) and *MOTHER OF FT* (*MFT*), etc.) were related to photoperiod and circadian clock pathways (Bouche et al. 2016). Ten genes (like *AGL19* and *SVP*, etc.) were in the vernalization and ambient temperature pathway, seven genes were involved in the Gibberellin Acid (GA) pathway, and nine genes are integrators or related to meristem response and developmental process. Four genes bound by AGL16 were not clearly defined for the flowering pathways (Zhao et al. 2011; Boxall et al. 2005; Xiao et al. 2009). It should be further noted that, besides GA, jasmonate acid (JA) signaling could also time flowering as well (Kazan and Manners 2013; Zhai et al. 2015; Wang et al. 2017; Bao et al. 2019). Genes in this pathway were directly targeted by AGL16 (Table S3), thus it’s possible that AGL16 modulates flowering time also through this pathway. Taken together, AGL16 might impact several flowering pathways, and the alteration of flowering time in mutants of *AGL16* could be a net effect of multiple flowering pathways.

**Fig 3.**
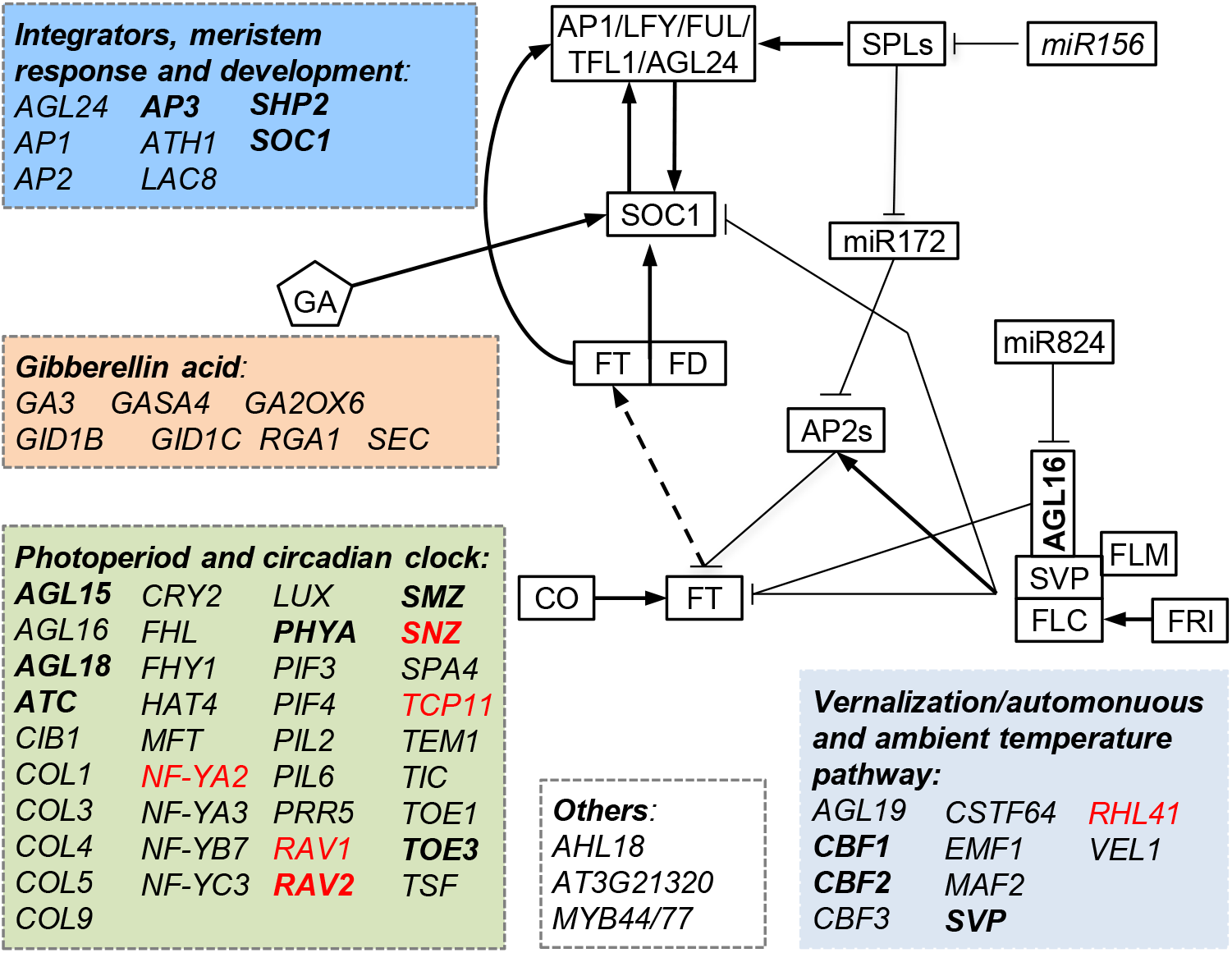
Molecular pathways (indicated with different color boxes) targeted by AGL16. Genes with names in bold were common targets for AGL16 and SOC1, while those in red were differentially expressed between the *agl16 soc1* and *soc1-2* mutants.

### AGL16 binds to *SOC1* promoter

The floral integrator gene *SOC1* was one of the targets bound by AGL16 (Fig. 4A; Table S3). AGL16 interacted with three DNA segments (peaks 1389, 1390 and 1391) in the promoter region of *SOC1* that harbored several CArG-motifs. Peak 1390 overlapped with a region bound by SOC1 itself (SOC1 binding region 1) (Tao et al. 2012), while peak 1389 overlapped with regions previously shown to be targeted by SVP (Tao et al. 2012; Mateos et al. 2015) or FLC (Deng et al. 2011; Mateos et al. 2015). An independent ChIP-qPCR assay confirmed AGL16 binding on all three peaks with the binding on peaks 1389 and 1391 relatively stronger than on peak 1390 (Fig. 5B). The second segment bound by SOC1 itself (SOC1 binding region 2 or fragment 7) was not targeted by AGL16. As AGL16 forms protein complexes with SVP and FLC (Hu et al. 2014), it is likely that AGL16 binds target regions together with these two MADS-box proteins, thereby participating in modulating target expression. However, the *SOC1* transcription was only weakly affected by loss-of-function of *AGL16* in the Col-0 background and not significantly in the Col-*FRI* background (Fig. 5C; Fig. S4), a pattern that had also been observed for *SVP* (Hu et al. 2014).

**Fig 4.**
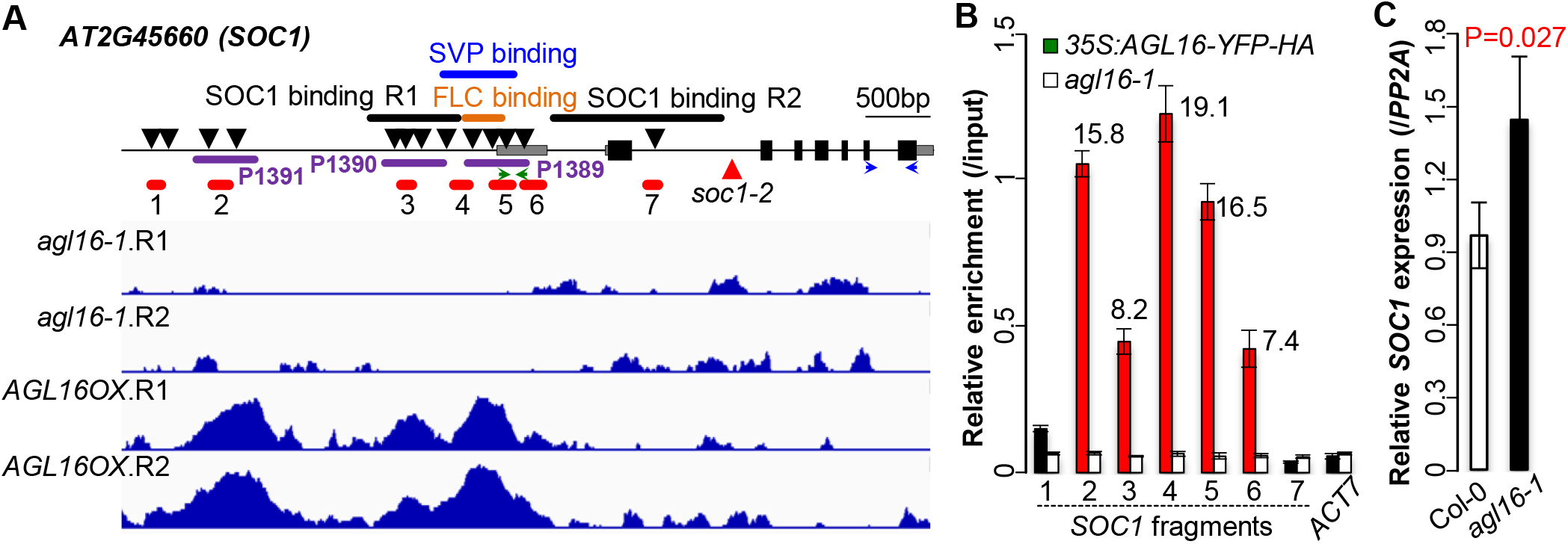
AGL16 targets and regulates *SOC1*. **A**. Schematic representation of the *SOC1* locus. Filled bars indicated exons and gray bars marked the 5’- and 3’-UTR regions while the line indicated the non-coding region of *SOC1*. Arrows downward labelled the putative CArG-boxes potentially bound by MADS-box proteins. The dark purple lines indicated the three peaks (P1389, P1390 and P1391) bound by AGL16. Orange, blue and black thick lines marked the known regions targeted by FLC, SVP and SOC1, respectively. Note that two sites in the regulatory region of *SOC1* were bound by itself (SOC1 binding R1 and R2; see ref. Tao et al. 2012). Red lines (1 to 7) showed the regions tested for AGL16-YFP-HA binding on *SOC1* chromatin. Horizontal arrows marked the position of primers used for quantification of 5’-UTR (green) and CDS (blue) regions. The lower panel showed the ChIP-seq profile at *SOC1*. **B**. Relative enrichment of AGL16 on *SOC1* chromatin tested with ChIP-qPCR. Fold change values with signficant enrichment was labelled above bars. *ACT7* was taken as a negative enrichment control. **C**. Relative expression of *SOC1* CDS against *PP2A* in Col-0 and *agl16-1* plants.

**Fig 5.**
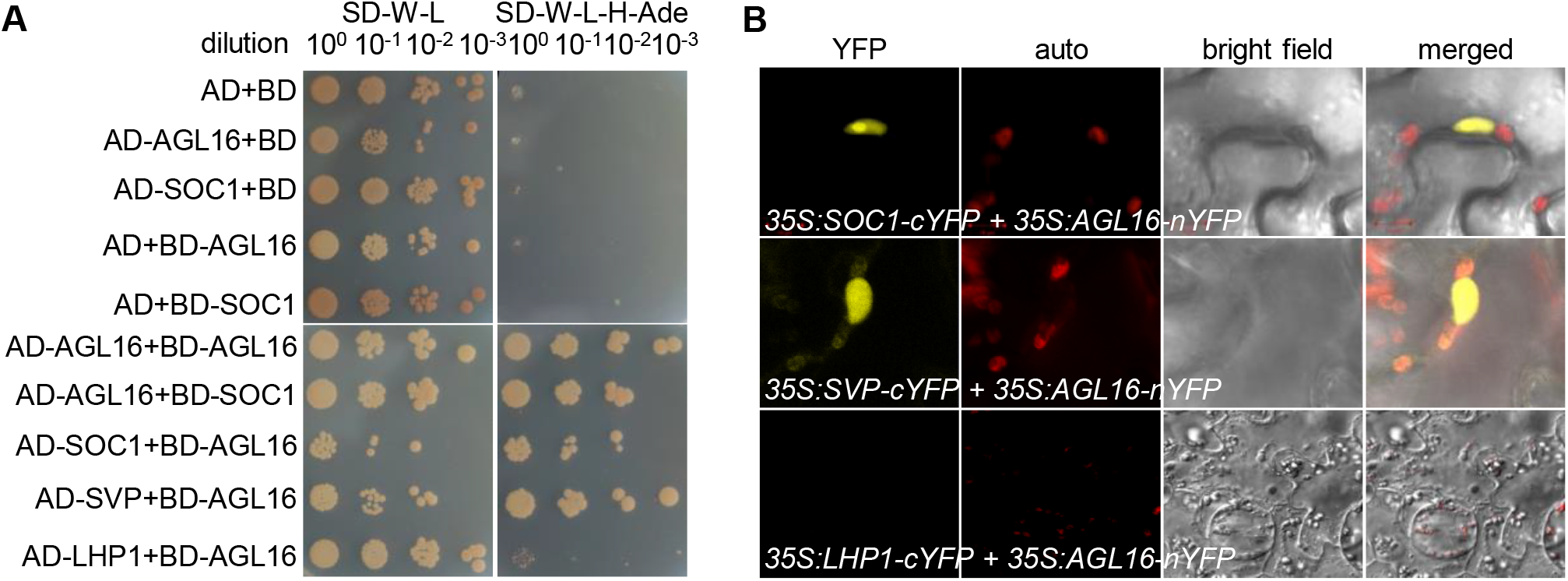
AGL16 forms protein complex with SOC1. **A**. Yeast two-hybrid assay revealed a direct interaction between AGL16 and SOC1. Each protein was fused to either the activation domain (AD) as prey or the DNA-binding domain (BD) as bait. Serial dilutions (10^0^ × to 10^−3^ x) of J69-4A cells containing different construct combinations indicated on the left were grown on control (left) and selective (right) medium. The AGL16-SVP and the AGL16-LHP1/empty vector combinations provided positive and negative controls, respectively. Note the formation of a AGL16 homodimer. **B**. BiFC assay evidenced the formation of AGL16-SOC1 complex in nucleus of *Nicotiana benthamiana* leaf epidermis. The interaction was tested with constructs *35S:SOC1-cYFP* and *35S:AGL16-nYFP*. A negative interaction between AGL16 and LHP1 and a positive interaction between AGL16 and SVP were tested as well (see also Hu et al. 2014).

### AGL16 can form protein complex with SOC1 and co-targets a common set of genes

Previously, AGL16 was been demonstrated to form heterodimer with SOC1 (Fig. S10) (de Folter et al. 2005; Immink et al. 2009). We verified this interaction with Yeast-2-Hybrid (Y2H) and bimolecular fluorescence complementation (BiFC) techniques. Y2H assays confirmed interactions between SOC1 and AGL16 (Fig. 5A), which was as strong as the previously reported direct interaction between AGL16 and SVP (Hu et al. 2014). LHP1 was used as a negative control. BiFC fusing the N-terminal half of yellow fluorescent protein (nYFP) with AGL16 (*35S:AGL16-nYFP*) and the C-terminal of YFP with SOC1 (*35S:SOC1-cYFP*) detected an interaction of AGL16 with SOC1 in the nuclei of Agrobacterium infiltrated tobacco leaves (Fig. 5B). Hence AGL16 and SOC1 can form complexes, which may contribute to the regulation of the expression of downstream targets.

We next examined whether AGL16 and SOC1 had common targets. For this aim, the previously generated binding profiles for SOC1 were used to identify shared targets with AGL16 (Immink et al. 2012; Tao et al. 2012). We applied the same annotation procedure for both AGL16 and SOC1 binding profiles in order to identify common genes. There were 193 AGL16 bound segments that overlapped with 240 SOC1 peaks (Table S4). These peaks were in the +/-2 Kb vicinity of 223 genes (five without annotation information), which were then taken as AGL16 and SOC1 common targets (Fig. 6A). Most of these common peaks were in the 1 kb region surrounding TSS with AGL16 peaks a bit more proximal (Fig. 6B). We further identified 211 CArG-box motifs in 144 common peaks (400 bp surrounding peak centers; 74.6% of all overlapped peaks) with MEME-ChIP. Eighty-seven peaks harbored one CArG-box (DCCAAAAAWGGAAAR; 60.4%), while the rest featured two (49 or 34%) or three (6 peaks or 4.2%) or more (2 peaks; Fig. S5A). The distances between the CArG-box motifs were significantly spaced with 20-40 bases (Fig. S5B). Among these common targets, genes involved in floral organ development (or reproductive growth) and responses to hormone stimulus including ethylene and ABA were significantly enriched (Fig. 6C; Table S4). Eight genes of the photoperiod and circadian clock related pathways (*AGL15, AGL18, ATC, PHYA*, *RAV2*, *SMZ*, *SNZ* and *TOE3*), three genes of the temperature-related pathways (*CBF1, CBF2* and *SVP*), and *SOC1* itself were involved in flowering (Fig. 3), indicating that AGL16 and SOC1 could act together to time floral transition in *Arabidopsis*.

**Fig 6.**
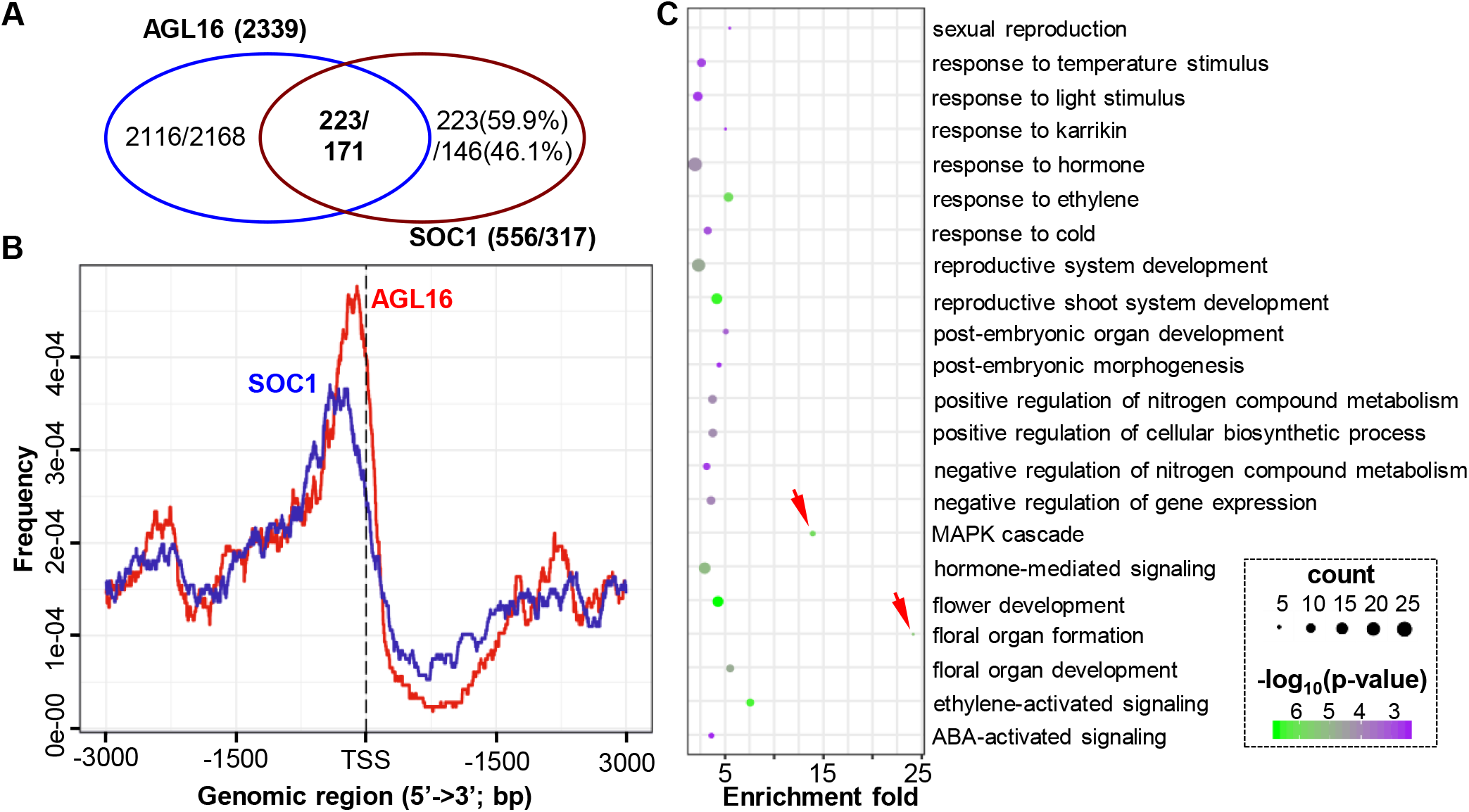
AGL16 and SOC1 share a common set of target genes involved in multiple functions. **A**. Venn diagram showing that 223/171 genes (Immink et al. 2012 / Tao et al. 2012) were co-bound potentially by both AGL16 and SOC1. **B**. Binding intensities for AGL16 (red) and SOC1 (blue) peaks surrounding transcription starting sites (TSS). Regions 3kb upstream and downstream of TSS were plotted. **C**. Selected significantly-enriched GO terms for the common targets. Note the GO terms marked by red arrowheads.

### The AGL16-SOC1 module is important for genome-wide gene expression and flowering time regulation

As AGL16 and SOC1 formed heteromeric protein complexes and as their genetic interaction played a role in the regulation of *AGL16* expression, we determined to what extent the gene expression at the genome-wide level could be affected by the AGL16-SOC1 module (Table S2). For this, we carried out a comparative transcriptomics analysis using the single and double mutants between the *agl16-1* and *soc1-2* lines. In contrast to the very broad binding spectrum of AGL16, we only detected very small number of genes showing differential expression (DEGs) in *agl16-1* single mutant (9 up and 12 down) compared to Col-0 (Fig. 7A; Table S5). The *soc1-2* single (155 up and 285 down) and the *agl16 soc1* double (49 up and 353 down) mutants had similar number of DEGs but *soc1-2* featured more up and less down DEGs *(Yate’s chi-square test*, p<0.001; Fig. 7A), indicating that AGL16 either countered SOC1’s repressive role on gene expression or its inductive role. A heatmap analysis of DEGs in the *soc1-2* vs Col-0 revealed that absence of *agl16* mostly reverted the differential gene expression observed in *soc1-2* to wild type levels (Fig. 7B). Genes down-regulated in the *agl16 soc1* mutants showed also down-regulation in *soc1-2* (Fig. 7C). In contrast, genes up-regulated in *agl16 soc1* were barely affected by either single mutation, suggesting that for these genes, AGL16 and SOC1 synergistically contribute to the repression. Accordingly, only 83 *soc1-2* DEGs (in total 155 up and 285 down; ~18.9%) overlapped with the *agl16 soc1* DEGs (375 up and 182 down; ~14.9%; Fig. 7D). Therefore, AGL16 has an important potential in regulating gene expression at the genome-wide level, but apparently depends on its genetic background, here, the *SOC1* activity.

**Fig 7.**
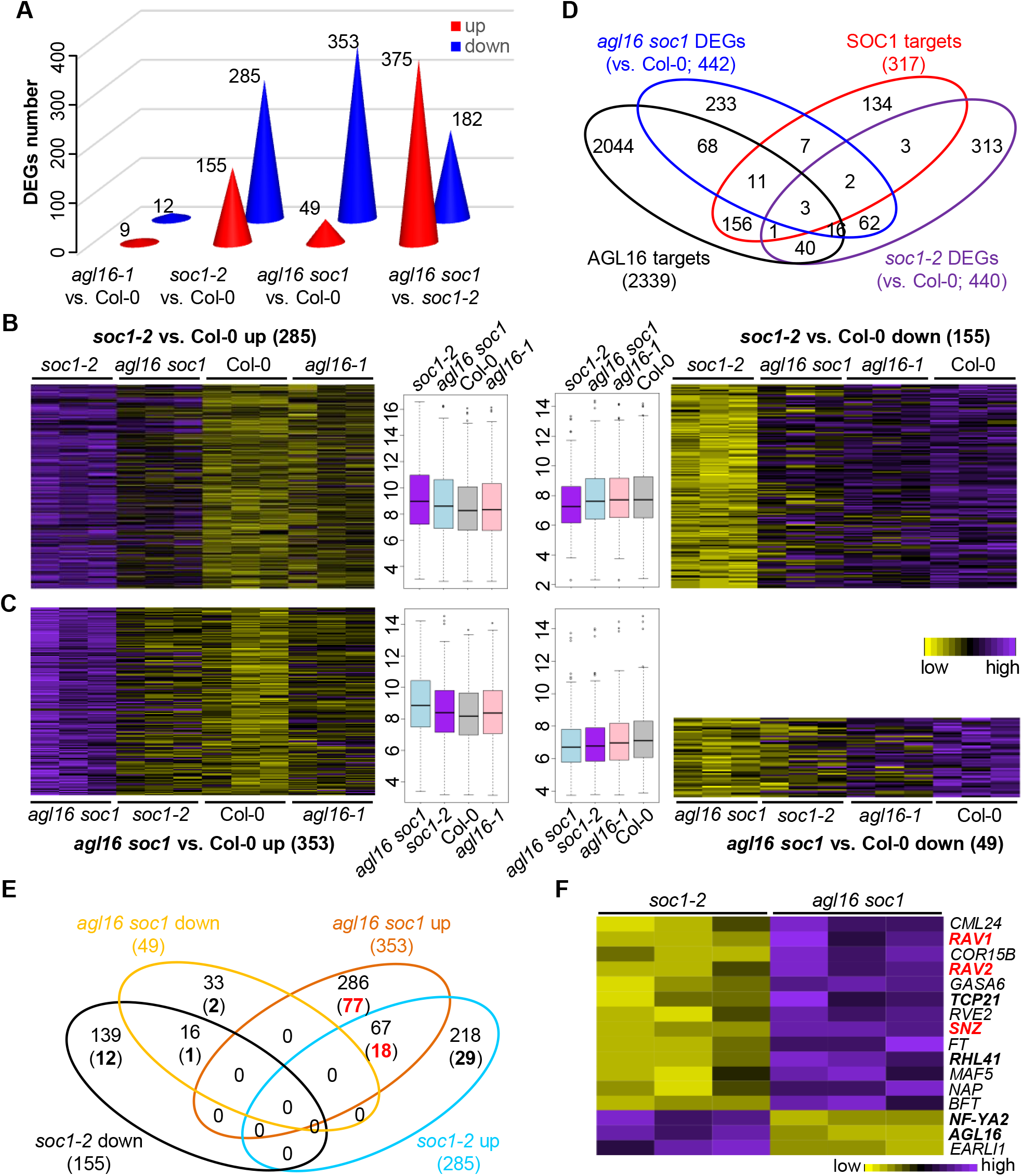
The *AGL16-SOC1* module collaborates on regulation of genome-wide gene expression. **A**. The number of differentially expressed genes (DEGs) in three mutants. The exact number of up (red) or down (blue) regulated DEGs were given on each cone. **B** and **C**. Heatmaps showing the normalized relative expression of *soc1-2* (**B**) and *agl16 soc1* (**C**) DEGs in all four lines. The boxplots in the middle gave the data distribution pattern for each cluster. **D**. Venn diagram demonstrating the overlap between DEGs and the AGL16 targets profile. **E**. A detailed comparison between the DEGs in soc1-2 and agl16 soc1 mutants with the AGL16 binding profile. Bold numbers in brackets showed the number of DEGs bound by AGL16. **F**. A heatmap showing the normalized relative expression of the DEGs related to flowering time regulation in the *soc1-2* and *agl16 soc1* mutants.

We next examined to what extent these DEGs associated with AGL16 targeting. Among the 557 *agl16 soc1* DEGs, AGL16 bound to 98 genes (~22.2%), of which only 23 (~4.1%) were also targeted by SOC1 (Fig. 7D). About 13.6% or 60 *soc1-2* DEGs were likely the AGL16 targets (Yate’s chi-square test, p=2e-8, in comparison to genome-wide level of AGL16 binding). However, we noticed that only nine *soc1-2* DEGs (~2% among 440) were potential targets of SOC1, a pattern similar to a previous report, in which 52 SOC1 targets were among the 1186 DEGs (Tao et al. 2012). There were six targets (~28.6%) showing differential expression in the 21 *agl16-1* DEGs.

Moreover, we identified more than a quarter of up-regulated DEGs specifically in the *agl16 soc1* (77 among 286) were AGL16 targets in contrast to about 13.3% of up-regulated DEGs specifically in the *soc1-2* mutant (29 among 218; Yate’s chi-square test, p=0.0035; Fig. 7E). Among the 67 up-regulated DEGs shared between the *soc1-2* and *agl16 soc1* mutants, 18 (26.9%) were potentially AGL16 targets. However, only less than 8% of down-regulated DEGs in both mutants were potentially targeted by AGL16. Together, these data suggest that AGL16 may act mainly as a transcriptional repressor in the *soc1-2* background.

Among the DEGs between *agl16 soc1* and *soc1-2* plants, we identified 17 known genes involved in flowering time regulation with seven of them (*NF-YA2, TCP2 1, RHL41, AGL16* and three *AP2-like* genes *RAV1, RAV2/TEM2*, and *SNZ*) being targeted by AGL16 (Fig. 7F; Table S5). Expression of *FT* was significantly enhanced in *agl16 soc1* double mutant. In line with this, the double mutant *agl16 soc1* flowered significantly earlier (~20 rosette leaves) than the *soc1-2* single mutant (~25.6 rosette leaves; about 21.6% reduction in rosette leaf number) but still later than both *agl16-1* (~11.1 rosette leaves; ~13.6% reduction) and wild type Col-0 (~12.9 rosette leaves) plants (Fig. 8). This indicated that *AGL16* could counteract *SOC1* effects in flowering time regulation. Thus, the regulatory role of *AGL16* in floral transition depends on *SOC1* function, similar to the genetic dependency of *AGL16* on *FLC* (Hu et al. 2014). It’s possible that SOC1 repressed the inhibition of AGL16 on *FT* expression, which should be tested further.

**Fig 8.**
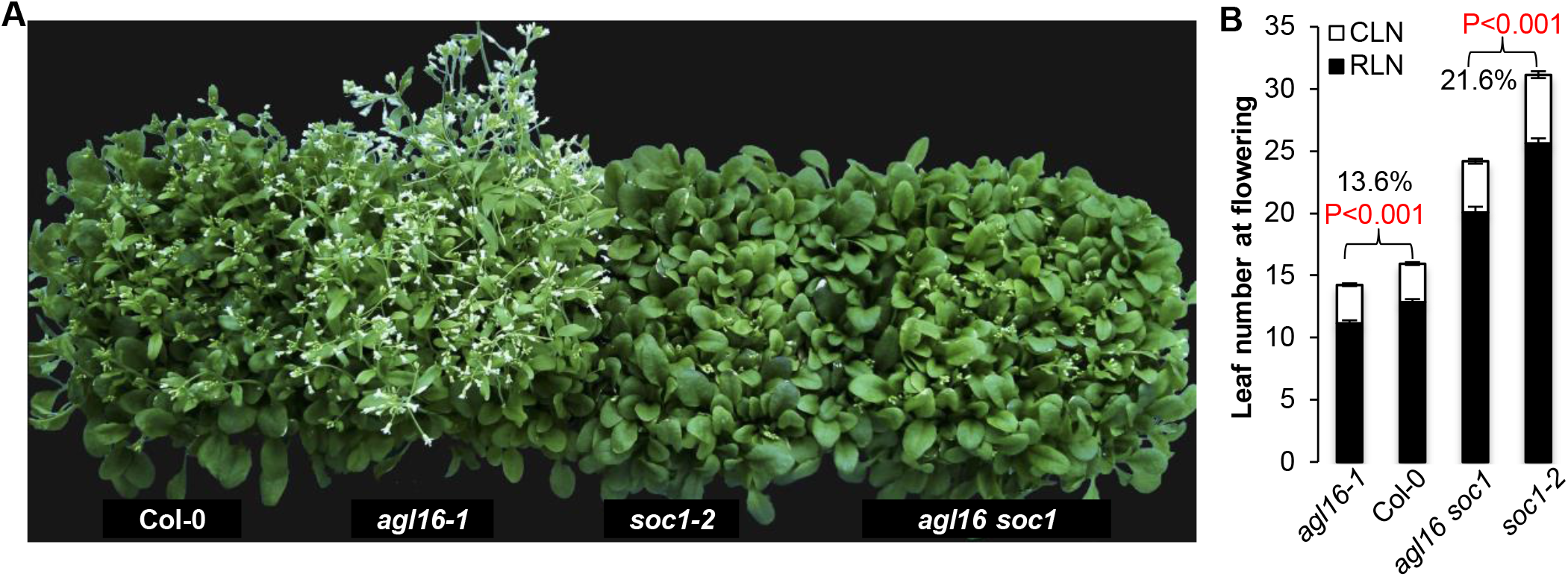
AGL16 and SOC1 regulate additively flowering time. **A**. Flowering behaviors of LD-growing wild type Col-0, *agl16-1, soc1-2* and *agl16 soc1* mutants. **B**. Leaf number production upon flowering under LD conditions. Rosette (filled bars, RLN) and cauline (open bars, CLN) leaves were shown. Numbers in percentage showed the earlier flowering level of *agl16-1* and *agl16 soc1* comparing to Col-0 and *soc1-2*, respectively. Analyses were repeated three times and all had similar patterns.

### *AGL16* is involved in water loss and seed germination regulation

Given the very broad binding profile at the genome-wide, we continued to explore whether AGL16 played a regulatory role in other biological process. AGL16 binds to a large set of genes involved in abscisic acid (ABA) signaling (29), and ABA (101) and water (62) responses (Table S3). Since the function of ABA in regulating adaptation to water availability has been well established, we questioned whether *AGL16* could have a role in water governance. We used the *agl16-1* and *m3*, a line in which the *AGL16*-specific negative regulator *miR824* was highly expressed (Hu et al. 2014; Kutter et al. 2007), to examine the water-loss-rate in the aerial parts of Arabidopsis plants after cutting. Compared to Col-0 control plants, six-weeks-old short-day grown mutant plants displayed a weak but significant decrease in water loss (2-4%; *Student’s t-test*, p<0.001; Fig. 9A), suggesting that *miR824*-regulated *AGL16* could regulate the response to water deficiency. The change in water loss could be either caused by the reduction of stomata density (Kutter et al. 2007) and/or by altering the stomata aperture size, which is tightly associated with the ABA signaling pathway (Zhao et al. 2020).

**Fig 9.**
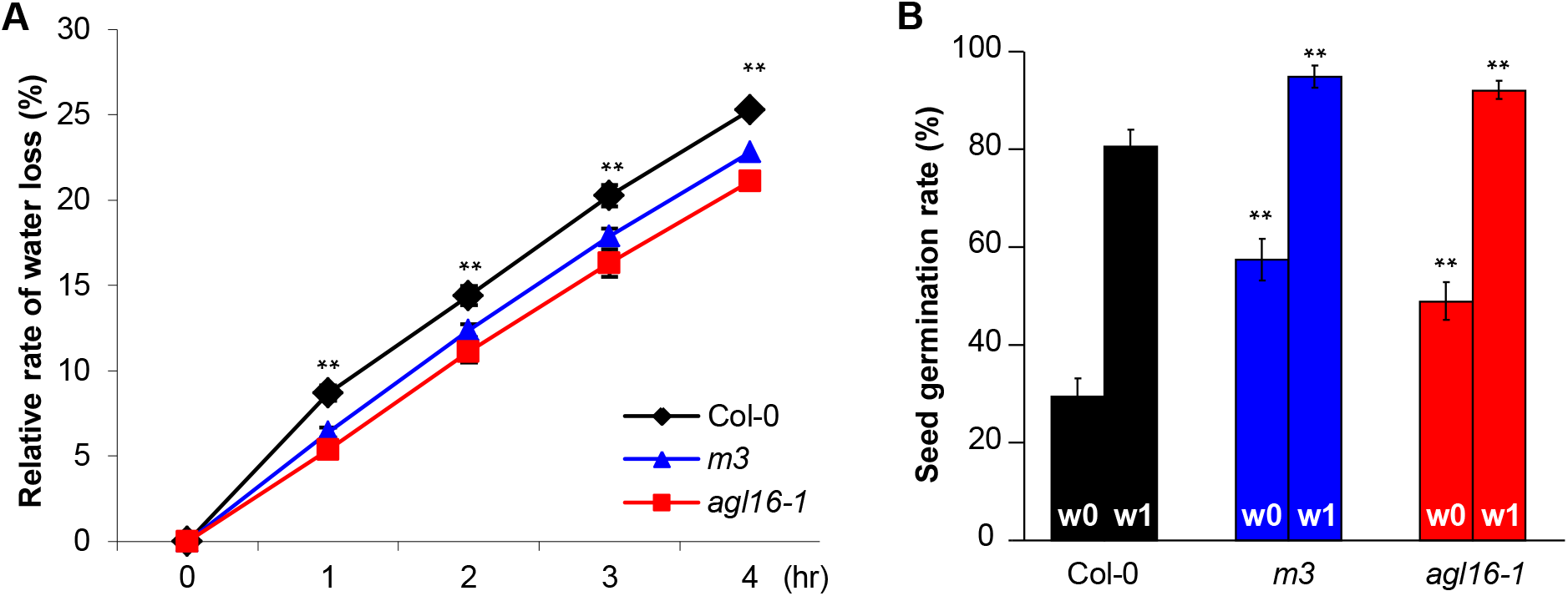
*miR824-AGL16* regulates water loss rate and seed dormancy. **A**. Relative rates of water loss for *agl16-1* and *m3*. Six weeks old rosettes growing under SD conditions were cut and the decreases of fresh weight in percentage were measured. **B**. Changes in seed germination behavior of *agl16-1* and *m3* lines in contrast to Col-0. Bars mark the germination proportion in percentage at the time point of freshly harvesting (w0) and one week after (w1). In A and B, mean values for at least ten individuals with standard deviation were shown. The experiments were replicated for at least twice with similar patterns. Significance was tested against Col-0 with *Student’s t-test*, ** p<0.01.

ABA plays essential roles in seed germination and dormancy control (Bewley 1997; Bentsink and Koornneef 2008). Not surprisingly, a further examination on seed dormancy levels demonstrated a significant alteration in germination rate of freshly harvested seeds of *agl16-1* and *m3* compared to the wild type, and to a reduced extent, after one-week storage (Fig. 9B). This pattern was tightly associated with increased levels of *miR824* and a decreased expression of *AGL16* in germinating seeds (Das et al. 2018). Taken together, these data suggest a regulatory role of AGL16 in water adaptation and seed germination. Corroborating with this, AGL16 was recently identified as a negative factor of drought resistance via regulation on stomata density and ABA accumulation (Zhao et al. 2020). *CYP707A3* (Zhao et al. 2020), *CYP707A1* and *AAO2*, which are involved in ABA biosynthesis and metabolism, were among the AGL16 targets (Table S3). Since ABA related signaling genes were also enriched in AGL16-SOC1 common targets (Fig. 6C), it would be worth to examine further the regulatory function of the AGL16-SOC1 module in water loss and seed dormancy processes.

## Discussion

In this study, via ChIP-seq and transcriptomic profiling as well as genetic analyses, we show that AGL16 targets to a broad range of genes and acts in a wide range of biological processes such as water deprivation and seed germination time. Depending on SOC1 function, AGL16 occupies important hubs in the GRNs involved in flowering time regulation.

### AGL16 interacts with SOC1 and times flowering with a partial background dependency on *SOC1*

AGL16 is known as a floral repressor in photoperiod pathway of flowering time regulation (Hu et al. 2014). Corroborating with previous notions (de Folter et al. 2005; Immink et al. 2012; Immink et al. 2009), AGL16 forms heteromeric protein complexes with SOC1, as evidenced by our Y2H and BiFC analyses (Fig. 5). This suggests that both proteins work together to target a common set of downstream genes. We provide evidences that AGL16 binds potentially more than 2000 target genes (Fig. 2), many of which are share with SOC1 (~50% of SOC1 bound genes; Fig. 6). Since AGL16 forms also protein complexes with SVP and FLC (Hu et al. 2014) and potentially with SEP3 (Fig. S3) (de Folter et al. 2005), it would be worth to examine whether AGL16 shares also common targets with these TFs. As *AGL16* times flowering time with a genetic background dependency on *SOC1*, similar to our previous finding on *AGL16*’s dependency on *FLC* activity (Hu et al. 2014), whether *SOC1* and *FLC* work together to mediate the AGL16’s function in flowering time regulation awaits further investigation.

AGL16 might exert its regulation potential in several pathways controlling flowering time (Fig. 3). Being congruent with its photoperiod dependency in regulation of flowering time, AGL16 targets 37 genes (including *AGL16* itself) related to photoperiod and circadian clock pathways. Under the tested environmental conditions *agl16-1* still shows a normal vernalization response (Hu et al. 2014), several genes related to temperature responses are directly targeted by AGL16. FLC, SVP and SOC1 might be partners of AGL16 in this respect as all three proteins target also directly on some of these temperature-related genes (Deng et al. 2011; Mateos et al. 2015; Immink et al. 2012; Tao et al. 2012). The binding of AGL16 may cause both positive and negative influences on the transcription of these targets (Fig. 7), which encompass both repressors and promoters of the floral transition. Indeed, several of the flowering time genes targeted by AGL16 show an enhanced or decreased expression when *AGL16* activity is modified in the *soc1-2* background (Fig. 3 and 7; Table S4, S5). Therefore, the early flowering phenotypes present in *AGL16* loss-of-function mutants (Fig. 8) (Hu et al. 2014) might be a net-effect/balance of the regulation on different pathways.

### *AGL16-SOC1* module acts in regulating genome-wide gene expression

MADS-box TFs often act together to target and regulate the expression of a broad set of downstream genes (de Folter et al. 2005; Deng et al. 2011; Immink et al. 2009; Immink et al. 2012; Kaufmann et al. 2009; Kaufmann et al. 2010; Lee et al. 2008; Mateos et al. 2015; Tao et al. 2012). Although AGL16 binds more than 2000 genes, which is in line with its very broad expression in many tissues and organs (Alvarez-Buylla et al. 2000), AGL16 alone can only affect the expression of a limited number of genes in the background of Col-0, in which *SOC1* is functional (Fig. 7). However, when *SOC1* is non-functional (in *soc1-2* background), AGL16 modulates the expression of more than 550 genes and acts both as a transcriptional repressor and activator. Moreover, in the *soc1-2* background, AGL16 seems mainly act as a transcriptional repressor as more than a quarter of the up-regulated DEGs, in contrast to the less than 8.5% of the down-regulated DEGs, are potential targets of AGL16. Hence AGL16’s activity in gene expression regulation requires partially the participation of SOC1. On the other hand, SOC1 also needs partially the *AGL16* function as SOC1’s repressive activity significantly drops (from 155 to 49 genes) but the promoting activity significantly increases (from 285 to 353 genes) when *AGL16* has no function. Many *soc1-2* DEGs are not differentially expressed any more in *agl16 soc1* mutant (Fig. 7). Therefore, AGL16 and SOC1 act both additively and synergistically in regulation of genome-wide gene expression.

### AGL16 is important in GRNs connecting life-history traits

Both AGL16 and SOC1 can directly bind to chromatin and regulate the expression of genes involved in hormone signaling and abiotic stresses (Fig. 6) (Immink et al. 2012; Tao et al. 2012). Corroborating with this, alteration in *AGL16* activity significantly changes the water loss efficiency, a process for which stomata development (Kutter et al. 2007) and ABA signaling might play a role (Zhao et al. 2020). Previously, SOC1 has been implicated in modulating stomata opening (Kimura et al. 2015). AGL16 also participates in the regulation of seed germination (Fig. 9), a key step in plant life cycle and adaptation to fluctuating environmental conditions (Koornneef, Bentsink, and Hilhorst 2002; Bewley 1997). The expression of *miR824*-regulated *AGL16* decreases significantly during seed germination (Das et al. 2018). The regulatory role of AGL16 in seed germination might be related to ABA as many ABA signaling genes including those encoding for ABA receptors, such as *PYL4, PYL5, PYL8*, are directly targeted by AGL16 (Table S3). *PYL8* is co-bound by SOC1 (Table S4). Therefore, the *miR824-AGL16* module seems to be important in GRNs connecting the two key transitional events, i.e., flowering and germination.

In summary, our data reveals that, as a master regulator in GRNs connecting multiple biological pathways, AGL16’s function depends partially on *SOC1*, similar to the genetic dependency on *FLC* (Hu et al. 2014). AGL16 might act as a glue, like other MADS-box TFs do, to modulate the chromatin accessibility of their interacting proteins to micro-tune the expression of downstream genes at proper stages and environmental conditions (Pajoro et al. 2014; Immink et al. 2009; Kaufmann et al. 2010; Richter et al. 2019). It will be important to address this further to understand their precise roles and mechanisms in balancing development and adaptation.

## Materials and methods

### Plant materials, and growth conditions

*A. thaliana* plants including wild-type Col-0, *agl16-1, 35S:AGL16-YFP-HA* in *agl16-1* background, Col-*FRI*, *agl16-1* Col-*FRI*, and *m3* have been described previously (Kutter et al. 2007; Hu et al. 2014). The *soc1-2* mutant in Col-0 background (Torti et al. 2012) was kindly provided by Prof. George Coupland. To test the genetic interactions between *AGL16* and *SOC1, agl16-1* and *soc1-2* were crossed and double mutant was screened with gene-specific primers (Table S1) (Torti et al. 2012; Kutter et al. 2007; Hu et al. 2014).

Arabidopsis seeds were stratified in distilled water at 4°C for 72 h and sown in soil and grown under LD conditions (16-h light at 21°C and 8-h night at 18°C). Seedlings for phenotyping were planted either in growth rooms or chambers, while materials for gene expression analysis and ChIP assays were sown on Murashige and Skoog medium plates (Hu et al. 2014).

### RNA Isolation, real-Time RT-qPCR, and RNA-seq assays

Total RNA was extracted with TRI Reagent®(Molecular Research Center, Inc. Cincinati, USA). Ten days old seedlings were used for quantification of relative expression of selected genes with *PP2A* as reference (Hu et al. 2014). Reverse transcription was carried out with the HiScript®II Q RT SuperMix for qPCR (+gDNA wiper) and quantification PCRs were performed with ChamQ™ SYBR qPCR Master Mix (both from Vazyme Biotech co. ltd, Nanjing) on QuantStudio™ 7 Flex Real-Time PCR System (ThermoFisher). Three to four biological replicates from each of two to three independent trials were applied for each experiment. A similar protocol was developed for monitoring relative enrichment of DNA fragments in ChIP-qPCR experiments. All the primers used in this study are included in Table S1.

For RNA-seq, materials were collected from three independent biological replicates for each genotype, and DNA-free total RNA was generated as described above. Illumina True-seq library preparation was performed from 3 μg DNA-free total RNA and sequenced by the Biomarker Technologies Corporation, Beijing, China. Quality trimmed pair-end RNA-seq reads were mapped to the Arabidopsis TAIR10 annotation using the *HISAT2* v2.1.0 (Kim et al. 2019). The *featureCounts* included in *subread* v1.6.4 package was applied to calculate reads counts on each gene (Liao, Smyth, and Shi 2013; Liao, Smyth, and Shi 2014). *DESeq2* v1.14.1 was used to detect differentially expressed genes (DEGs; fold change above 1.5 and p.adj<0.1). Only uniquely mapped reads were used for downstream analysis. Transcriptional clustering analysis was performed using the *heatmap.2* function in *R*. GO analysis was performed with *PANTHER* in TAIR web-tool (https://www.arabidopsis.org/tools/go_term_enrichment.jsp) (Mi et al. 2017) or *agriGO* pipeline (Tian et al. 2017).

### ChIP-seq and ChIP-qPCR assays and data analysis

ChIP experiments were carried out following protocols described (Zhou et al. 2016; Reimer and Turck 2010). Chromatin for both *agl16-1* and *agl16-1 AGL16OX* plants was extracted from ten-day-old seedlings grown under LD conditions at ZT14, and precipitated with antibody against GFP (Abcam, Ab290). For ChIP-seq, the immuno-precipitations from two independent trials were used for NGS library preparation with NEBNext®Ultra™ II DNA Library

Prep Kit for Illumina®(E7645, New England BioLabs Inc.) and high-throughput sequencing with HiSeq2000 platform. ChIP-seq reads were mapped to the TAIR10 assembly of *A. thaliana* using *BWA-MEM* (v0.7.17-r1188) (Li 2013). Reads with mapping quality below 30 were discarded with *SAMtools* v1.7 (Li et al. 2009). Duplicated reads were removed using *Picard MarkDuplicates* v1.119. The resulted .*bam* file was used as input to call AGL16 enriched regions with MACS v2.2.7.1 (Zhang et al. 2008). Enriched regions were generated by the comparison of immune-precipitated products to input for *AGL16OX* and then compared against *agl16-1*. For annotation of AGL16 targets, the *R* package *ChIPseeker* was used (Yu, Wang, and He 2015). The position and strand information of nearest genes were reported with the distance from peak to the TSS of its closest gene identified. As annotations might overlap, we use ‘promoter’ definition in *ChIPseeker* as the highest priority for annotation. Each binding site was assigned to only one gene. IGV was used for data visualization of the binding profiles for targets (Thorvaldsdottir, Robinson, and Mesirov 2013). Enriched motifs in AGL16 binding peaks were identified using *Homer* suite with *findMotifsGenome.pl* function (Heinz et al. 2010). Motifs in AGL16-SOC1 co-targeted regions were analyzed with *MEME-ChIP* tools (Machanick and Bailey 2011), and the spacing between primary and secondary motifs was analyzed with *SpaMo* (*spamo-dumpseqs-bin 20-verbosity 1-oc spamo_out_1 -bgfile./background -keepprimary -primary DCCAAAAAWGGAAAR*). We compared the AGL16 targets to SOC1 targets from both Immink (2012) and Tao (2012) with the same annotation procedures for AGL16 (Immink et al. 2012; Tao et al. 2012). In an earlier independent trial, we pooled the immune-precipitations from two biological replicates and sequenced the products. This pooled sequencing results gave similar pattern of AGL16 targets profile but with a lower coverage hence the data was not shown. *Yate’s chi-square tests* were performed online (http://www.quantpsy.org/chisq/chisq.htm). The ~400 flowering time genes were downloaded from https://www.mpipz.mpg.de (Bouche et al. 2016) with self-curations. Reads data for RNA-seq and ChIP-seq experiments were accessible at NCBI under accession code SUB5067038.

### Phenotype assays

Flowering time assays were carried out according to previous report (Hu et al. 2014). Four independent trials were applied and each gave similar pattern. Phenotype comparisons were performed with Student’s *t-test* with *Bonferroni-correction*.

For water-loss assays, rosette leaves of six week-old plants grown under short day conditions were used for measurements of water-loss rates (Lefebvre et al. 2006). Fresh rosettes were cut at their base and immediately weighted to establish initial fresh weight (FW_i_). These rosettes were left in open air at room temperature in the lab and weighted 1, 2, 3, and 4 hrs after cut to calculate weight loss per unit of time, (FW_t_-FW_i_). At each time point, the amount of water lost was quantified by expressing the lost weight per unit of time as a percentage of FW_i_. To quantify the role of the *miR824/AGL16* regulatory system in the rate of water loss, average water loss of the mutants was expressed as percentage of the average water loss measured in Col-0. This experiment was repeated two times and both gave similar pattern.

For seed dormancy assays, about 50 individually- and freshly-harvested seeds were plated onto a filter paper moistened with demineralized water in Petri dishes and incubated in LD conditions in transparent moisturized containers (16h light/8h dark, 25°C/20°C cycle) (Xiang et al. 2014). Germination was scored after 7 days of incubation. For each assay, at least three trials, each with minimum 10 individual plants, were used.

### Yeast two-hybrid and biomolecular fluorescence complementation (BiFC) experiments

Yeast two-hybrid and the BiFC assays were carried out to test the physical interaction between AGL16 and SOC1 proteins according to previous report (Hu et al. 2014). In yeast two-hybrid assay, interactions between AGL16-SVP and AGL16-AGL16 were applied as positive controls while the AGL16-LHP1, AGL16-BD, SOC1-BD, AD-AGL16, and AD-SOC1 were applied as negative controls together with empty vectors. For BiFC assay in *Nicotiana benthamiana* plants, *35S:SOC1-cYFP* construct was built by cloning the full-length encoding-region without stop codon of *SOC1* (from Col-0) into *pDONR221* entry vector first and later transferred into *RfA-sYFPc-pBatTL-B* vector. The interactions between AGL16 and SVP, between AGL16 and LHP1, were used as positive and negative controls, respectively.

## Acknowledgements

We thank Liangyu Liu, Feihong Yan, Fei He, Yibo Sun, Shulan Chen for assistance in experiments. This work was supported by grants from National Natural Science Foundation of China (31570311 to J-Y H, 31501034 to X D, 31700275 to Y-X D, 31800261 to F C), from the CAS Pioneer Hundred Talents Program (292015312D11035 to J-Y H), and CAS Key Laboratory for Plant Diversity and Biogeography of East Asia to J-Y H, from the China Postdoctoral Science Foundation (2017M613023 to Y-X D), the Postdoctoral targeted funding from Yunnan Province to F C and Y-Y D, and the Yunnan basic and applied research funding to F C. This work is partially facilitated by the Germplasm Bank of Wild Species of China. The authors declare no conflict of interest.

## Author Contributions

J-Y H conceptualized and coordinated the research; L-P Z performed the ChIP experiments and collected the RNA samples, D-M Y and Y Z carried out the protein interaction assays, X D, J-Y H and L-P Z created the genetic materials and did the genetic analyses, X D analyzed and visualized the data, F C, Y-X D, X-D J, F-M Q and F T did other analyses; J-Y H wrote the paper with help from F T and the other authors. All authors had read and approved the manuscript.

Supporting information

